# Ghosts of symbionts past: The hidden history of the dynamic association between filarial nematodes and their *Wolbachia* endosymbionts

**DOI:** 10.1101/2025.03.17.643741

**Authors:** Emmelien Vancaester, Guy R. Oldrieve, Alex Reid, Georgios Koutsovoulos, Dominik R. Laetsch, Benjamin L. Makepeace, Vincent Tanya, Sven Poppert, Jürgen Krücken, Adrian Wolstenholme, Mark Blaxter

**Author notes:** Emmelien Vancaester: Georgios Koutsovoulos: Institute of Computer Science (ICS), Foundation for Research and Technology - Hellas (FORTH), Nikolaou Plastira 100, Vassilika Vouton, GR-70013 Heraklion, Crete, Greece Dominik R. Laetsch: FRAMIDOSUR S.L., Av. TT.OO. Neckermann 24, 35100 Maspalomas, Gran Canaria, Spain.

## Abstract

Many, but not all, parasitic filarial nematodes (Onchocercidae) carry intracellular, maternally-transmitted *Wolbachia* symbionts, and these alphaproteobacteria are targets for anti-filarial chemotherapeutic interventions for human disease. The symbionts of Onchocercidae derive from four of the major supergroups (C, D, F and J) defined within the genus *Wolbachia*. Using twenty-two whole genome sequences of filarial nematodes and genome sequences of their *Wolbachia* partners, we have explored the evolutionary history of the nematode-*Wolbachia* symbiosis. We screened the nuclear genome sequences of all the nematodes for nuclear *Wolbachia* transfers (NUWTs), fragments of the *Wolbachia* genome that have been integrated into the nuclear genome.

Six species have no current *Wolbachia* infection. *Setaria labiatopapillosa* had no validated NUWTs and we interpret this to mean that this species was never infected with *Wolbachia*. In the other five species (*Acanthocheilonema viteae, Cercopithifilaria (Ce.) johnstoni, Elaeophora elaphi, Loa (Lo.) loa* and *Onchocerca flexuosa*) we found NUWTs, implying they have previously had and have now lost *Wolbachia* infections. For each NUWT locus, we identified the supergroup membership of the *Wolbachia* from which it originated, and found that the five *Wolbachia*-free species carried NUWTs derived from multiple supergroups, including a high shared rate of sequences derived from supergroup C. In *Dirofilaria repens* we identified a sample that carried two *Wolbachia* symbionts, one from supergroup C and one from supergroup F. In *Dirofilaria immitis* where live infection with a supergroup C *Wolbachia* is found, we identified NUWTs derived from an F *Wolbachia*, confirming that the F association predated the divergence of these *Dirofilaria* species. The supergroup D lineage of *Wolbachia*, as present in the human parasites *Wuchereria bancrofti* and *Brugia malayi*, derives from a replacement event. *Madathamugadia (Md.) hiepei* shows signs of multiple recent repeated endosymbiont replacement.

From these data we infer that the history of *Wolbachia* in onchocercid nematodes includes not only cospeciation (as present in the *Onchocerca-Dirofilaria* group in association with supergroup C *Wolbachia*) and loss (in the five *Wolbachia*-free species), but also frequent symbiont replacement and dual infection. This dynamic pattern is challenging to models that assume host-symbiont mutualism.

## INTRODUCTION

*Wolbachia* are bacteria belonging to the Rickettsiales (alpha-proteobacteria), a group characterised by their intracellular lifestyle and wide range of phenotypic effects on their hosts [1,2]. *Wolbachia* is an endosymbiont of terrestrial arthropods and parasitic nematodes [3]. In most arthropods *Wolbachia* endosymbionts are non-essential and hosts can be cured of infection by treatment with antibiotics. Arthropod *Wolbachia* generally promote their own transmission by reproductive manipulation, whereby female hosts carrying infections have higher Darwinian fitness than uninfected females [2]. In several insects it has been shown that *Wolbachia* infection can also be beneficial by promoting host resistance to viral or protozoan parasite infection. *Wolbachia* may thus play a role in controlling disease transmission [4,5]. *Wolbachia* of haematophagous bed bugs provide B-vitamins to their hosts [6]. *Wolbachia* infection of nematodes is usually portrayed as more mutualist than parasitic, as experimental elimination of the endosymbiont with antibiotics also harms the nematode [7]. It has not been possible to generate experimental evidence of reproductive manipulation in nematode-*Wolbachia* interactions because all individuals of infected species carry the endosymbiont.

*Wolbachia* is classified into supergroups based on analysis of molecular markers. Supergroups A and B are the most common and are restricted to terrestrial arthropods (largely in the Insecta). Most other supergroups also infect terrestrial arthropods, but often have restricted host ranges. For example, supergroups I and V are found in fleas [8], supergroup P in quill mites [9] and supergroup S in pseudoscorpions [10]. *Wolbachia* have been described from plant- and animal-parasitic nematodes. Some *Pratylenchus* plant-parasitic nematodes are infected with supergroup L *Wolbachia* [11,12]. The tylenchoid nematode *Howardula* sp., which infects sphaerocerid flies, was shown to contain *Wolbachia* with an unusually small genome, assigned to supergroup W [13]. Most *Wolbachia* infections have been described in the family Onchocercidae, or filarial nematodes. Filarial nematodes are parasites of vertebrates, including humans, and many have been shown to carry *Wolbachia* [14,15]. The filarial nematode *Wolbachia* are from supergroups C, D, F and J [15,16]. Interestingly, *Wolbachia* supergroup F also occurs in arthropods.

Lateral transfers of *Wolbachia* DNA into the host nuclear genome (nuclear-*Wolbachia* transfers, or NUWTs) have been identified frequently in filarial nematodes [17–21] and in arthropods [21–26]. Insertions have been observed in the genomes of individuals not currently infected in both nematodes [27] and arthropods [28]. These “fossil” NUWTs provide evidence of past infection. NUWT insertions range in length from a few hundred base pairs to near-full *Wolbachia* genomes [21,25,29]. There is no strong evidence that insertions are functional, and most display evidence of disabling mutations and pseudogenisation [30].

While early analyses suggested extensive co-phylogeny between *Wolbachia* in supergroups C and D with their nematode hosts [31], subsequent wider analyses identified uninfected species nested within infected clades, and breaks in co-phylogeny [15]. To explore the phylogenetic history of the dynamics of host-symbiont interaction we developed a toolkit to robustly identify NUWTs in host genomes. Here, we use this “fossil record” of NUWTs to explore the dynamics of *Wolbachia* association with these nematode hosts. We identify a surprisingly mobile symbiome, with multiple instances of endosymbiont loss and endosymbiont replacement.

## RESULTS

### Nematode genome phylogeny

We collated genome data from twenty-two onchocercid nematode species (Table 1). We present new whole genome assemblies for *Setaria labiatopapillosa*, a filarial parasite of cattle that is an outgroup to the other species sampled, and *Acanthocheilonema viteae,* a rodent parasite. We also generated additional data for *Dirofilaria (Dr.) repens*. We annotated the genomes and identified 670 single-copy orthologues. Using these we re-estimated the relationships of the Onchocercidae. The phylogeny is fully resolved with maximal support for every node (Figure 1A). Rooting the phylogeny with *S. labiatopapillosa*, we recapitulate the phylogeny produced by Lefoulon *et al* [32] based on seven loci. We included representatives of four of the five clades of Onchocercidae analysed by Lefoulon *et al.*, lacking only a representative of their ONC1 group, which arises basally to *S. labiatopapillosa*. *Elaeophora elaphi* was not included in the analyses of Lefoulon *et al.*, but its position, as sister to *A. viteae* and *Litomosoides (Li.) sigmodontis*, is as expected [33].

**Table 1:**
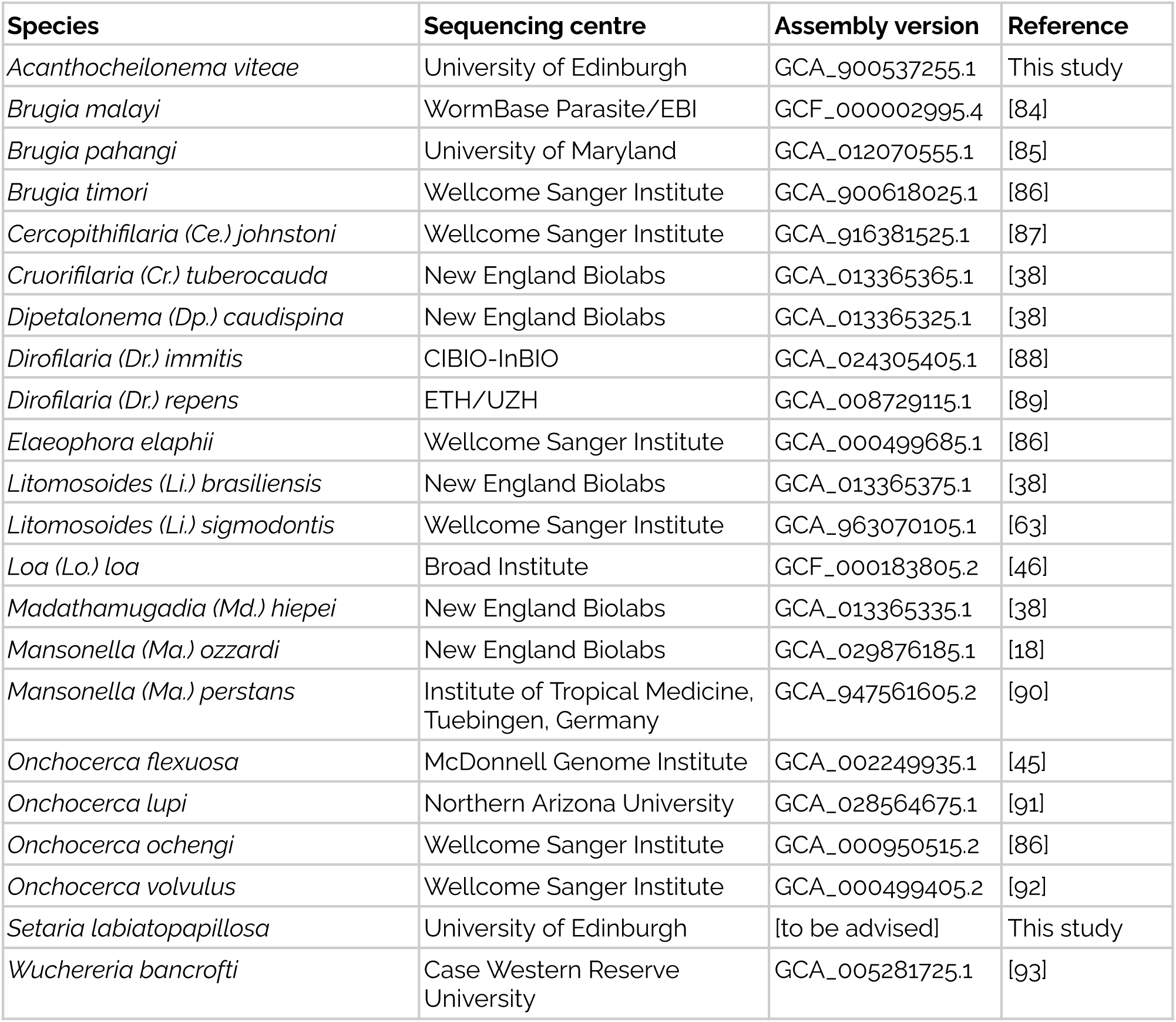
Nematode genome data

**Figure 1:**
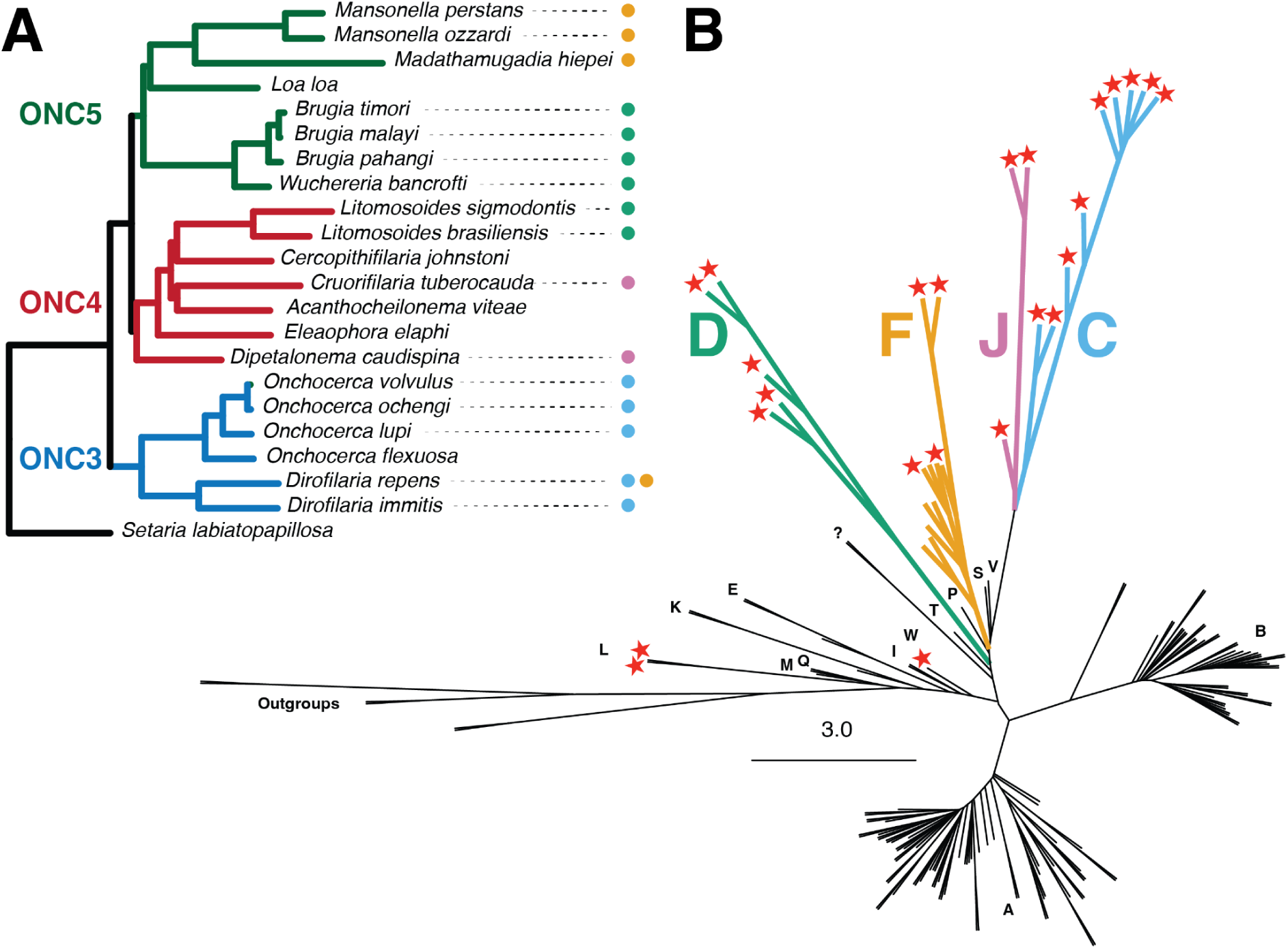
Genome phylogeny of filarial nematodes and *Wolbachia*. **A** Phylogeny of twenty-two filarial nematodes (Onchocercidae) based on a concatenated alignment of 670 nuclear-encoded, orthologous proteins, analysed with IQ-TREE. All nodes gained maximal support (ML bootstrap = 100%). The Onchocercinae subtaxa defined by LeFoulon *et al.* [32] are indicated (ONC3, ONC4, ONC5). For each nematode species, current infection with *Wolbachia* is shown by a coloured dot, and the supergroup allocation of the endosymbiont is indicated by colours as in part B. **B** Unrooted phylogeny of 167 *Wolbachia* genomes based on analysis of 696 orthologous proteins with IQ-TREE. Letters A-W indicate assigned supergroups. Red stars indicate *Wolbachia* from nematode hosts. Support for deep branching between supergroups, and within supergroups C, D, F and J, is near-maximal. See Supplementary Figure S2 for a full representation of the phylogeny with tips named.

### Wolbachia genomes

We used BlobToolKit [34] to identify *Wolbachia* symbiont genomes in newly sequenced nematode genomes. No symbiont genomes were detected in *S. labiatopapillosa* or *A. viteae*. In whole genome data from two *Dr. repens* individuals, we identified variable presence of *Wolbachia*. One sample contained a *Wolbachia* from the expected supergroup C. The other contained the supergroup C genome, and an additional, larger supergroup F genome with distinct coverage and GC content (Figure S1) (Figure 1B). Both *Wolbachia* assemblies were deemed complete based on the Rickettsiales dataset within CheckM, at 98.1% (supergroup C genome) and 97.6% (supergroup F genome) completeness.

We collated genome data from 1,444 isolates of *Wolbachia* (Table S1), from 242 host species, including *Wolbachia* genomes for all the *Wolbachia*-infected nematodes analysed in this study (apart from *Brugia timori* and *Onchocerca lupi*, where *Wolbachia* assemblies were not available). To avoid potential biases due to different annotation protocols for published *Wolbachia* genomes, we re-predicted proteomes for all strains with prokka/prodigal [35]. Using orthoFinder [36], we generated an orthology clustering for the 1,444 newly-predicted proteomes from 16 different supergroups (A-F, I-M, P, S, T, V, W) and 7 outgroup species (see Figure S2). A large fraction of the orthogroups were singletons (6,659 proteins) or strain-specific orthogroups (144 orthogroups).

The majority (97.5%) of the 1,667,683 proteins were clustered into 1,386 multi-genome orthogroups. Only 432 orthogroups had members from all 16 *Wolbachia* supergroups, but 725 had representatives from all of the well-represented supergroups A-F. A large set of 2,386 orthogroups was found to be specific to a *Wolbachia* supergroup. Most of these were specific to supergroup D (841), followed by A (519), C (316), B (265), K (197), F (88), M (52), E (44), V (28), I (20), L (9) and J (7). Two hundred families were only present in supergroups A and B but were missing in all others, and are mainly composed of transposases.

Rarefaction analyses, using only genomes with high completeness, showed major differences between *Wolbachia* supergroups in terms of their predicted size of their pan-proteome, the total protein diversity likely to be present in all strains of each supergroup. For *Wolbachia* as a whole, the rarefaction curve levels off after addition of the proteomes of about 30 strains, suggesting that the proteome diversity of *Wolbachia* likely includes 3,000 orthogroups (Figure S3). Within supergroups A and B, diversity appeared to asymptote at approximately 2,200 orthogroups after the addition of around 30 proteomes (Figure S3A). The C, D and F pan-proteomes did not reach asymptote, likely because too few proteomes were available, but indicated pan-proteome sizes of 1,300 orthogroups in supergroup F, 1,370 orthogroups in supergroup C and 1,290 orthogroups in supergroup D (Figure S3B).

To reduce redundancy across the *Wolbachia* genomes, a set of 165 strains retaining maximal sequence diversity in the dataset was selected using the dRep toolkit [37] (Table S2). We derived a robust *Wolbachia* phylogeny using a concatenated protein supermatrix derived from 696 one-to-one orthologues present in at least 75% of the selected strains (Figure 1B). This phylogeny supported the distinctiveness of the different supergroups. Between-strain divergences within supergroups A and B were relatively small compared to the divergence observed within supergroups C, D, J and, to a lesser degree, F. The supergroup topology was similar to previously described phylogenies. The nematode-infecting *Wolbachia* were interspersed with minor arthropod-infecting clades. The nematode-infecting supergroups C and J were sisters. Supergroup F is sister to a clade comprising supergroups C, J, S and V, and supergroup D is sister to a clade of C, J, V, S, P and S *Wolbachia*. This structure reflects that described previously [13,38,39]. The supergroup F *Wolbachia* infecting *Dr. repens* had a slightly unexpected position within the supergroup, as it clustered closest to genomes of the insect-infecting supergroup F *Wolbachia* rather than other nematode-infecting supergroup F genomes (Figure 1B). A closely related supergroup F strain in the nematode *Cercopithifilaria (Ce.) japonica* was recovered among *Wolbachia* infecting the bed bug *Cimex lectularius* and termites from the genus *Coptotermes* and *Odontotermes* [15,16].

### NUWTs in twenty-two filarial nematode genomes

NUWTs are an instructive marker of historical association between a nuclear genome and *Wolbachia* [27]. Even if *Wolbachia* infection has been cleared, NUWTs can persist as genomic fossils in the nuclear genome. As the vast majority of NUWTs are non-functional in their new host genome [30], they evolve neutrally with no constraint on small insertions and deletions, and thus can be hard to detect by simple sequence similarity search. We used our *Wolbachia* protein orthogroups to develop tools to identify NUWTs in the filarial nematode nuclear genomes.

For each of the 2,647 *Wolbachia* orthologue clusters containing more than three members across three different strains, we generated a protein alignment [40], and back-translated each alignment into the original DNA sequences [41]. While direct sequence- or sequence profile-based searches are sensitive to changes in frame that result from insertion and deletion mutations, hidden Markov models (HMM) are able to identify similar sequences even in the presence of insertions and deletions [42]. We derived a HMM from each DNA alignment and used these HMMs to screen the nematode nuclear genomes for NUWTs [43]. Almost half of the HMMs (1,179, 46%) identified putative NUWTs in the filarial genomes at an E-value cutoff of 1e-30. Using IQ-TREE [44], we generated phylogenetic trees of the *Wolbachia* genes and putative NUWT sequences from all 1,179 clusters that had putative NUWTs. These phylogenies were rooted and then parsed to discriminate between true NUWTs, with unequivocal signal of similarity to *Wolbachia* genes, and nematode genomic fragments identified due to a lack of specificity in the HMMs.

We identified a total of 7,906 initial matches grouped in just over 5,000 regions with between 12 matches in 12 regions (*S. labiatopapillosa*) and 818 matches in 653 regions (*Onchocerca volvulus*) in each nematode genome (see Table S4). For most matches, the filarial nematode nuclear genome-derived sequences nested within the diversity of sequences from *Wolbachia* (see Supplementary Figures S4-S8 for examples of these phylogenies). The 12 HMM hits to 7 orthologous families identified in *S. labiatopapillosa* all had higher sequence similarity to sequences from the other filarial species analysed and to other non-filarial nematode species than to *Wolbachia* sequences, and thus likely derive from false-positive, off-target matches to nematode nuclear sequences. We interpret these data as indicating that *S. labiatopapillosa* has no evidence of current or ancient infection with *Wolbachia* [15,32].

This low level of false positives gives us confidence in interpreting NUWTs identified in the nuclear genomes of other *Wolbachia*-negative filaria. It may be that our ability to identify NUWTs is negatively impacted by low genome quality or over-purging of fragments that have similarity to *Wolbachia*. However, we found similar patterns of NUWT abundance in closely related species despite the differences in assembly technologies and contiguity. The impact of NUWTs not observed because of technical issues will be minimal, and at worst we will have missed true associations rather than inferred false associations.

### Congruence and incongruence in the origins of NUWTs

Placing the supergroup origin of live *Wolbachia* infections on the nematode phylogeny, it is evident that the current pattern of presence across the filaria requires multiple infections with symbionts, including at least two distinct instances of infection with supergroup J and supergroup D strains and five independent losses of infection (Figure 1A). Exploration of the distribution of NUWTs indicates that the hidden history of association between *Wolbachia* and filaria is even more complex (Figure 3).

Several patterns were evident in the frequency and origins of NUWTs in onchocercid genomes. Firstly, nematodes infected with supergroup C *Wolbachia* had more NUWTs than those infected with supergroup D (Figure 2). In *Dirofilaria* and *Onchocerca* species carrying live supergroup C *Wolbachia* (i.e., *Dr. immitis, Dr. repens, O. volvulus*, *O. ochengi*, and *O. lupi*) we found between 483 and 653 NUWT regions, while in *Brugia* species, *W. bancrofti,* and *Litomosoides* species infected with supergroup D *Wolbachia* there were between 24 and 247. However, the inserted regions in *Brugia* species were on average larger. NUWTs of D origin in *Brugia* species were on average 2,038 bp, while NUWTs of C origin in *Onchocerca* species were on average 434 bp. The total inserted NUWT length was more than 400 kb in *Brugia* species, but only 188 kb to 306 kb in *Onchocerca* species (Figure 2B). The longest NUWT region detected, in *Brugia malayi*, was 29 kb.

**Figure 2:**
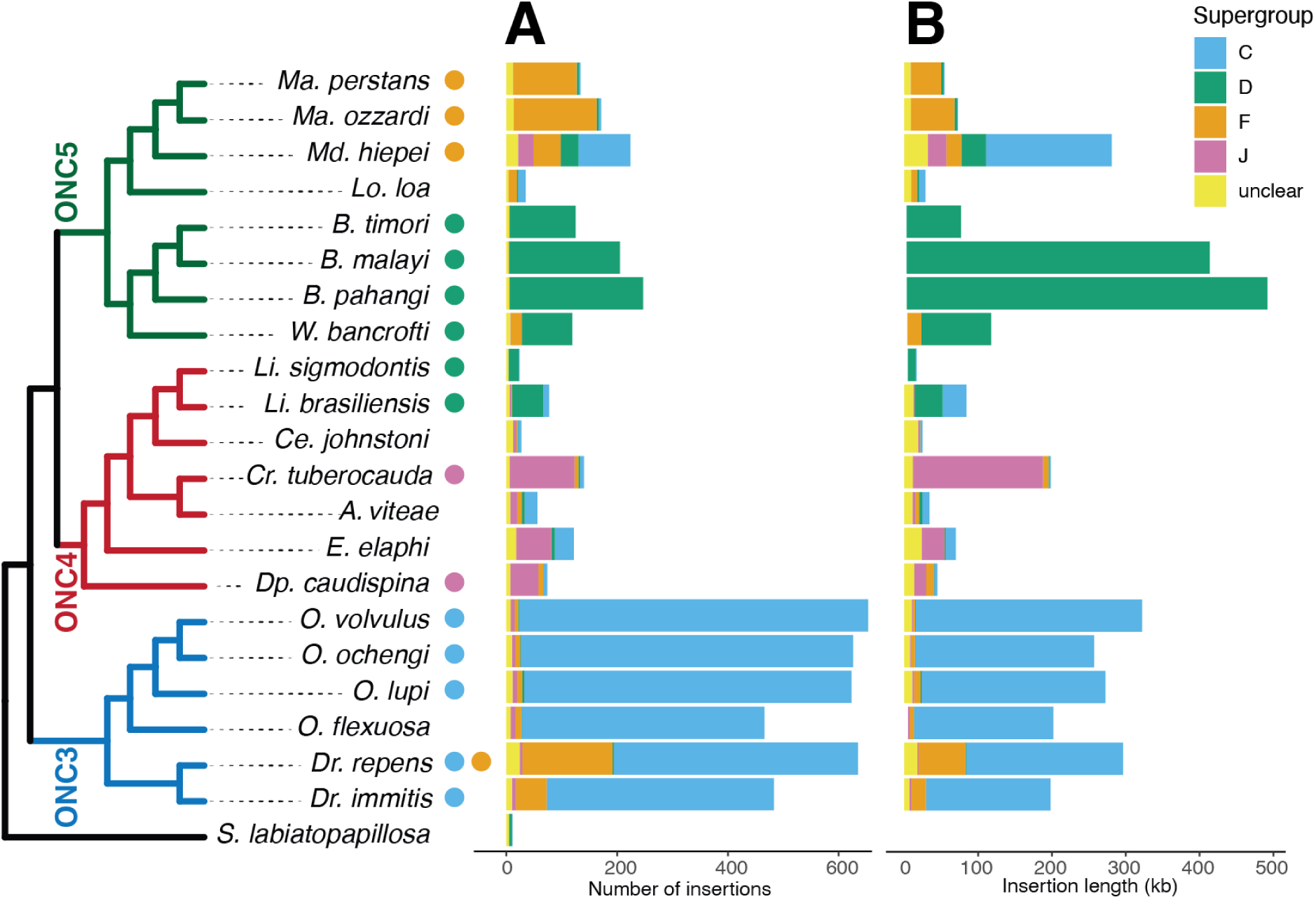
NUWTs in filarial nematode genomes. **A.** For each nematode species tested, the stacked histogram shows the classification by supergroup of origin of the NUWTs identified by Hidden Markov Model searching. NUWTs that had no clear phylogenetic association with any sequence from live *Wolbachia* are labeled “unclear”. Data are available in Supplementary Table 3. **B.** The sum of the lengths of the *Wolbachia* insertions for each species, colour coded by supergroup of origin. The cladogram to the left shows the relationships of the nematodes (from Figure 1A), and the coloured dots show the current infections of each species.

The NUWTs found in a species with a live *Wolbachia* infection were largely identified as having come from the same supergroup as the live infection. For example, 96% of the NUWTs in *O. volvulus* were derived from supergroup C *Wolbachia*. The majority of the remainder was assigned to supergroups F and J (one percent each). Whether these non-C classifications are real or due to unavoidable phylogenetic noise, perhaps due to short sequences evolving neutrally, is unclear. This pattern was especially strong in *Brugia* species, with 95-98% of NUWTs deriving from supergroup D *Wolbachia*, and the remainder largely unclassified. NUWTs derived from supergroup C genomes were observed in all filarial nematodes except for the *Brugia* and *Wuchereria* clade.

The *Wolbachia*-free nematodes *A. viteae, Ce. johnstoni, E. elaphi*, *Lo. loa* and *O. flexuosa* all carried NUWTs, attesting to their having once been infected. NUWTs have been described previously in *O. flexuosa* [19,45] and *Lo. loa* [46]. These filarial nematode species that have lost their *Wolbachia* infections (i.e., are aposymbiotic) mostly had fewer insertions with lower span than related species that have retained *Wolbachia* infection. We identified only 466 NUWTs in *O. flexuosa* compared to the >600 found in the other *Onchocerca* spp.. As the ancestor of *O. flexuosa* that eliminated its *Wolbachia* infection might be expected to have had the same number of NUWTs as other *Onchocerca* do today, this suggests that *O. flexuosa* has been losing NUWTs since it became aposymbiotic or that other *Onchocerca* species have had longer to accumulate NUWTs. The *Wolbachia*-free *E. elaphi* had more NUWT regions (122) than the closely-related, *Dipetalonema (Dp.) caudispina*, which is infected by a supergroup J *Wolbachia* (74). *A. viteae, Ce. johnstoni* and *Lo. loa* had only 56, 27 and 35 NUWTs, respectively. If NUWTs are lost and gained in a clock-like fashion, this implies that *A. viteae, Ce. johnstoni* and *Lo. loa* lost their *Wolbachia* earlier than did *E. elaphi*. The NUWTs carried by *Wolbachia*-free nematodes were frequently derived from different supergroups than currently found infecting their closest relatives. For example, *Lo. loa* is sister to the supergroup F *Wolbachia*-infected *Mansonella-Madathaumugadia* clade in ONC5, but carries both supergroup F and supergroup C NUWTs. Similarly, *A. viteae* and *E. elaphi* carry supergroup C-derived NUWTs as well as the supergroup J NUWTs expected from supergroup J infection of their phylogenetic neighbours *Dp. caudispina* and *Cr. tubercauda*.

### Dynamics of NUWT evolution show *Wolbachia* superinfection and replacement

In *Dr. repens*, there were 635 NUWT regions, 69% from C and 26% from F sources. Given the identification of a supergroup F *Wolbachia* in one of the two isolates sequenced, we interpret this difference in frequency as having arisen from the continuous presence of C *Wolbachia* in the species, and lower rates or duration of infection with F *Wolbachia*. In *Dr. immitis*, while most (85%) of the 483 NUWTs were derived from supergroup C *Wolbachia*, 58 (12%) were derived from supergroup F. For example, *Dr. immitis* carried two C-derived and one F-derived NUWTs derived from horizontal transfer of orthogroup OG0000236 (dihydrolipoyl dehydrogenase) genes, which also identified *Onchocerca* C-derived and *Md. hiepei* and *Dp. caudispinia* J-derived NUWTs (Figure S4). This suggests that *Dr. immitis*, or perhaps the common ancestor of the genus *Dirofilaria*, was once infected by a supergroup F strain, or that the species is currently variably infected by an F strain that has not yet been sampled. In this context, the F-like NUWTs in *O. volvulus* are intriguing as they may point to ancient or current rare infection with an F supergroup *Wolbachia*. We did not find any F supergroup NUWT regions that were likely to be orthologous between *Onchocerca* and *Dirofilaria*.

We found evidence for repeated *Wolbachia* replacement in the genome of *Md. hiepei.* Significant fractions of *Md. heipei* NUWTs were classified as deriving from all of the filarial nematode-infecting supergroups (42% from supergroup C, 22% from F, 14% from D and 12.5% from J). Fifty-eight out of the 349 orthogroups that had *Md. heipei* NUWTs contained multiple sequences from *Md. heipei* that likely originated from different *Wolbachia* supergroups. Eight contained NUWTs assigned to three different supergroups, e.g. OG0000277 (peptide deformylase; Supplementary Figure S5).

We identified many cases of sets of NUWTs that clustered in phylogenetic trees and recapitulated the phylogenies of their hosts. Most of these groups of related NUWTs were found in the four closely related *Onchocerca* species or the three *Brugia* species. Some supergroup C-derived NUWT groups included all *Onchocerca* members, suggesting they were inserted in the last common ancestor of the *Onchocerca* clade (e.g. OG0000301; Supplementary Figure S6). Similar patterns were observed for many D supergroup NUWTs in *Brugia* species (e.g. OG0000453; Supplementary Figure S7). As previously noted in the strongyloidean nematode *Dictyocaulus viviparus*, NUWTs can be duplicated within their host genomes [27]. We identified several NUWT groups that were present in multiple copies in their host genomes, particularly in the genomes of *Brugia* spp. (e.g. ATP-dependent zinc metalloprotease FtsH OG0000206; Supplementary Figure S8).

We explored whether NUWT integration in the genome is biased towards regions enriched in certain features. We overlaid repeat annotation with NUWT integration sites to detect enrichment towards certain repetitive classes and detected strong enrichment for the long terminal repeats from the family Pao. This was significant in all *Brugia* and *Dirofilaria* genomes, *O. volvulus*, *O. flexuosa*, *A. viteae*, *Cr. tuberocauda*, *W. bancrofti*, *Dp. caudispina* and *E. elaphi*. Furthermore, we found a strong association between helitrons and NUWT integration specific to the genus *Onchocerca*, *O. volvulus*, *O. flexuosa*, *O. lupi* and *O. ochengi.* No association with repeats was identified in *Md. hiepei*, the two *Mansonella* species, *Ce. johnstoni, O. ochengi* and *O. lupi*.

## DISCUSSION

We used nucleotide HMM built from genes present in living *Wolbachia* to detect fossils of *Wolbachia* sequence inserted into the genomes of filarial nematodes. This NUWT identification pipeline appears to be an honest estimator of association between nematode nuclear genomes and *Wolbachia* symbionts. The pipeline will not detect NUWTs that derive from *Wolbachia* genes that were singletons in our orthology clustering, or from sequences that are not part of protein-coding genes. It will also fail to distinguish *Wolbachia*-derived sequence from other nematode non-coding sequence when sufficient time and thus mutational change has happened. The pipeline will therefore tend to underestimate the true extent of NUWT DNA in the nuclear genome. The pipeline also relies on the assumption that the HMM are specific to *Wolbachia* genes, and that spurious matches to nuclear genome sequences are not inflating the count of hits. Based on its phylogenetic position, *S. labiatopapillosa* is believed not to have had an association with *Wolbachia* [15,32]. The potential NUWTs identified in *S. labiatopapillosa* were small, and grouped with other likely nematode nuclear genome-resident sequences. We thus believe that the method generates few false positives.

The lack of NUWTs in *S. labiatopapillosa* is in keeping with the model of LeFoulon *et al.* [15,32] that the relationship between *Wolbachia* and filarial nematodes was established in the ancestor of groups ONC3 (*Dirofilaria* and *Onchocerca*), ONC4 (*Dipetalonema*, *Acanthocheilonema*, *Cruorifilaria*, *Cercopithifilaria* and *Litomosoides*) and ONC5 (*Wuchereria*, *Brugia*, *Loa*, *Madathamugadia* and *Mansonella*) (Figure 1A). This parasitic group has been estimated to have diverged at the start of the Paleogene around 60 million years ago, a period of rapid mammal diversification [47].

The subsequent history of *Wolbachia* infection in these filarial nematodes is complex, and here we propose a model for this set of associations (Figure 3). Almost all of the analysed genomes, even those from species that lack current live infection with *Wolbachia*, have supergroup C NUWTs. This suggests that the original infection was with a supergroup C (or C-like) *Wolbachia*, which left NUWTs in the genomes of all descendant species (Figure 3). All analysed species in ONC3 that have a live infection with a supergroup C *Wolbachia* have a high level of supergroup C NUWTs. *O. flexuosa* has secondarily lost its *Wolbachia* infection [20] and has a lower load of supergroup C insertions than its close sister species, possibly because of ongoing stochastic loss and/or sequence drift. Many NUWTs in the genomes of *Onchocerca* species were orthologous (Supplementary Figure S6) and must have originated from integration in the genome of a common ancestor.

**Figure 3:**
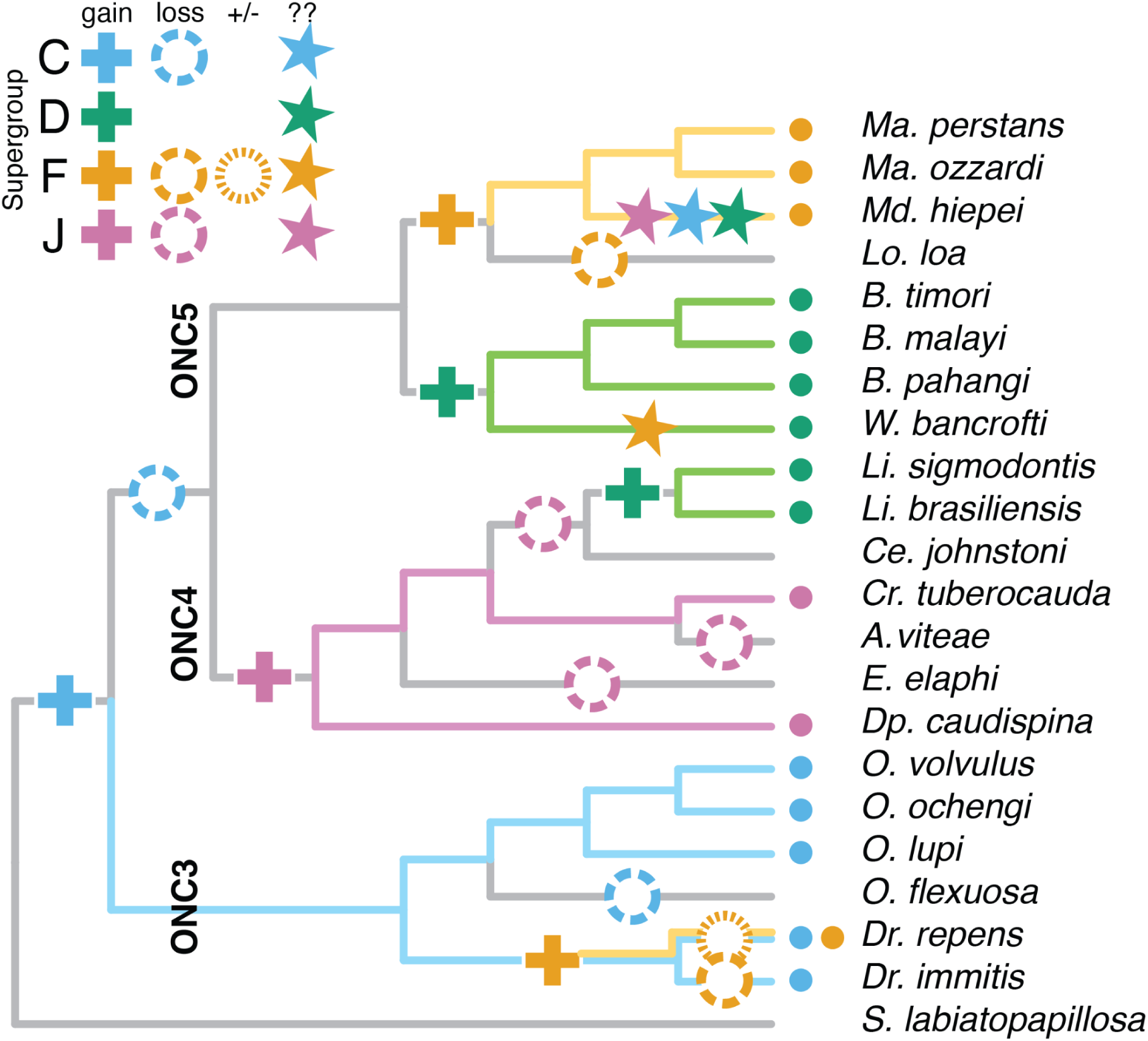
Origins of NUWTs in symbiotic and aposymbiotic filarial nematodes. Hypothesis of *Wolbachia* association through filarial nematode evolution. The cladogram shows the relationships of the nematodes (from Figure 1A), and the coloured dots show the current infections of each species. Plus signs indicate acquisition of *Wolbachia*, with the supergroup indicated by colour. Coarse-dashed circles indicate loss. The variably present supergroup F *Wolbachia* in *Dirofilaria repens* is shown with a fine-dashed circle. Stars indicate species with an unexpected frequency of NUWTs from *Wolbachia* supergroups that may indicate hit-and-run associations.

All genomes of nematode group ONC3 have NUWTs derived from supergroup F *Wolbachia*. Both *Dirofilaria* genomes have an elevated number of insertions of supergroup F-derived NUWTs (12 to 25% of all NUWTs). We found that *Dr. repens* can be simultaneously coinfected with supergroup C and supergroup F *Wolbachia*. While coinfection, especially involving supergroup A and B strains, is quite common across arthropod hosts [26,48,49], this is to the best of our knowledge the first demonstration of dual infection in nematodes. The potential for dual infection is supported by the demonstration, based on PCR amplicons of bacterial small subunit RNA loci (16S) in distinct specimens of the related species *Dirofilaria hongkongensis*, of supergroup C and supergroup F *Wolbachia* [50]. We suggest that the dual infection observed in *Dr. repens* is likely to have been present in the common ancestor of *Dr. immitis*, *Dr. repens*, and *Dr. hongkongensis* and this was the source of the frequent supergroup F NUWTs in *Di immitis*. Supergroup F infection has either been secondarily lost in *Dr. immitis* or is present in an as-yet-unsampled subset of individuals of this species. (Figure 3).

NUWTs assigned to a supergroup F source were identified in nearly all genomes, with only *E. elaphi* and *Li. sigmodontis* having none. While it is possible that assignment of some NUWTs to supergroup F is the result of noise in the phylogenetic reconstructions, in many cases the allocation of the NUWT to supergroup F was strongly supported. Supergroup F *Wolbachia* are remarkable in being the only supergroup known to infect both arthropods and nematodes, and the phylogeny of F genomes has shown that there are two distinct lineages of nematode-infecting F *Wolbachia* [16]. The newly sequenced F genome from *Dr. repens* clustered together with the supergroup F genomes from *Mansonella*, suggesting limited co-phylogeny with filarial nematodes. It is striking that NUWTs in the strongyloidean nematode *Dictyocaulus viviparus* were attributable to a supergroup F *Wolbachia*, despite no occurrence of a live infection in this or any other strongyle nematode being identified to date [27]. The lungworms (Strongyloidea) are not closely related to filarial nematodes and are estimated to have diverged 300 to 400 million years ago [47]. If the NUWTs in *D. viviparus* are the result of a “hit-and-run” infection of limited phylogenetic perdurance, the scattering of supergroup F NUWTs across the filaria could also result from similar brief “hit-and-run” infections with supergroup F *Wolbachia*.

NUWTs derived from supergroup D were common in *Brugia* and *W. bancrofti* in ONC5 and in *Litomosoides* species in ONC4. This pattern suggests two independent acquisitions of the supergroup D *Wolbachia*. The supergroup D NUWTs in *Brugia* were the longest detected in this analysis, with both the longest NUWT and the longest average NUWT size (Figure 2B). These may reflect recent insertions that have not yet been fragmented by neutral genomic processes. The other clade in ONC5, containing *Mansonella* species, *Md. heipei* and *Lo. loa*, has been infected by supergroup F *Wolbachia*. *Md. heipei* is currently infected by a supergroup F *Wolbachia*, but carries significant numbers of NUWTs for supergroups C (42% of NUWTs), D (14%), F (22%) and J (12%). This atypical NUWT distribution (Figure 2) suggests that this species or its ancestors has experienced multiple “hit-and-run” infections in its relatively recent evolutionary history.

The phylogenetic relationships of nematodes in ONC4 suggest that an ancestral species was infected with supergroup J *Wolbachia*, retained in *Cr. tubercauda* and *Dp. caudispina*. This *Wolbachia* has been lost in *A. viteae* and *E. elaphi* and has subsequently been replaced by a supergroup D *Wolbachia* in the *Litomosoides* species. In both *Li. sigmodontis* and *Li. brasiliensis* we found NUWTs derived from supergroup C *Wolbachia* (2 and 11 regions, respectively), and in *Li. brasiliensis* these accounted for about one-third of the span of the NUWTs identified. These supergroup C-derived NUWTs may point to historical infection by a C *Wolbachia* in these species or a *Litomosoides* ancestor.

Overall, we have shown that NUWT analysis allows us to assess the evolutionary history of interaction between *Wolbachia* and its hosts. It was originally proposed that *Wolbachia* show strong co-phylogeny with their filarial nematode hosts with exclusively vertical transmission [31], as opposed to supergroups A and B where host-switching events among arthropods are rampant [39]. The dynamic nature of the association between *Wolbachia* and filarial nematodes we infer here implies that there is unlikely to be a single explanation for the association [51], and it remains open whether the associations are mutualist (through metabolic supplementation, or through manipulation of vector or mammalian host immunity), parasitic reproductive manipulation (like the A and B supergroup *Wolbachia* in arthropods), or “ransom” or “addiction” based (where loss of live *Wolbachia* in germline cells results in toxin-mediated necrosis [52,53], and thus impacts host fitness [54]). The frequent discovery of *Wolbachia* in onchocercine nematodes, and the lack of evidence of infection in many other nematode groups, may be due to chance, a particular genetic susceptibility to infection in filaria, or their intimate relationship with *Wolbachia*-infected arthropods. This susceptibility may in turn have led to filarial nematodes being a fertile evolutionary playground for competing *Wolbachia* supergroup lineages.

We discovered horizontal transfers of DNA between symbionts and their hosts through a library of nucleotide HMMs representing symbiont genes that were able to identify transferred fragments, even if they had been subject to substitution, insertion and deletion. This HMM approach is equally applicable to finding NUWTs in other host groups and, given the building of HMM libraries for genes from representative symbiont genomes, insertions derived from other prokaryotic and eukaryotic symbiont taxa. We expect that, as in this nematode-*Wolbachia* example, hidden histories of symbiosis will emerge from deployment of such tools.

## Author contributions

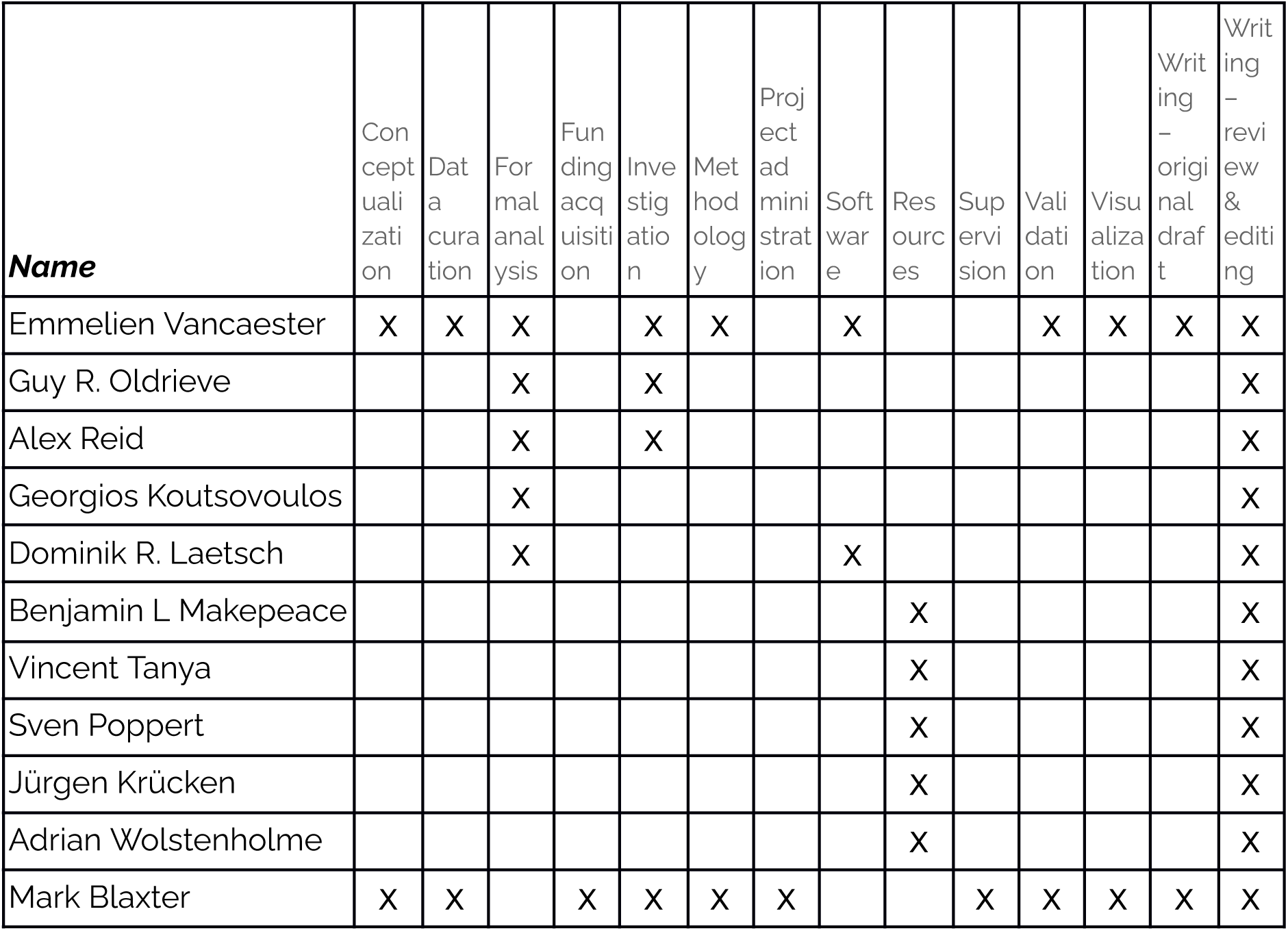

## ACKNOWLEDGEMENTS

This research was funded by the Wellcome Trust Grants 206194 and 218328. For the purpose of Open Access, the author has applied a CC BY public copyright licence to any Author Accepted Manuscript version arising from this submission. We thank Lewis Stevens and Marcela Ulianio da Silva for comments on the manuscript. We gratefully acknowledge the technical assistance of David Ekale in recovering *S. labiatopapillosa* specimens from Ngaoundéré abattoir, Cameroon. Genomic data for *Dr. repens* and *S. labiatopapillosa* were generated by the skilled team at Edinburgh Genomics (https://genomics.ed.ac.uk/).

## MATERIALS AND METHODS

### Sequencing and assembly of Setaria labiatopapillosa and Acanthocheilonema viteae genomes

A single cryopreserved specimen of *S. labiatopapillosa* was supplied by co-author Vincent Tanya from zebu (*Bos indicus*) tissue obtained from Ngaoundéré abattoir, Adamawa Region, Cameroon. After pulverisation of the specimen, DNA was extracted using a standard proteinase K digestion, followed by phenol-chloroform extraction and precipitation using isopropanol. DNA was pelleted by centrifugation and suspended in extraction buffer (EB; 10 mM Tris.HCl ph 8). An *A. viteae* DNA sample was provided by Kenneth Pfarr from the University of Bonn in Germany.

One genomic library of approximately 300 bp insert size was prepared for each species by Edinburgh Genomics, and sequenced (paired end, 100 bases) on Illumina HiSeq 2500. Sequence fastq files were submitted to the Short Read Archive under bioprojects PRJEB7555 (*S. labiatopapillosa*) and PRJEB1697 (*A. viteae*). The quality of Illumina reads was checked with FASTQC (http://www.bioinformatics.babraham.ac.uk/projects/fastqc). Correction was carried out using BLESS [55]. A preliminary single end assembly was screened for potential contaminating host sequences with BlobToolKit [34]. Production assemblies were carried out using SPAdes [56] (version 3.12.0). The assemblies were scaffolded with SCUBAT2 (https://github.com/GDKO/SCUBAT2) using cognate RNA-Seq data. Contigs less than 500 bp were discarded.

### Identification and assembly of two strains of *Wolbachia* from *Dirofilaria repens*

DNA extracts from two *Dr. repens* adult nematodes were analysed. One was from an experimentally infected dog infected with an isolate from Italy and the other was from a human who had lived in Croatia before the worm was detected (samples AW and S8 from Yilmaz *et al.* (2016) [57]; the ethics statement concerning the sampling of these nematodes is available therein). A genomic library of approximately 300 bp insert size was prepared by Edinburgh Genomics, and sequenced (paired end, 100 bases) on Illumina HiSeq 2500. Low-quality bases and adapter sequences were removed using fastp [58] (version 0.12.3) (cut_by_quality, cut_window_size 4, cut_mean_quality 20). A draft assembly was generated using SPAdes (version 3.12.0). BlobToolKit analysis (version 1.0) revealed the presence of 3.04 Mb of scaffolds that could be assigned to *Wolbachia* based on sequence identity. These scaffolds were grouped into two clusters with distinct GC content and read coverage (Figure S1). Reads were mapped to both bins and the two genomes assembled using megahit (version 1.2.9) [59] with complete tRNA sets (34 and 33 tRNAs coding for all amino acids).

### Data collation

We collated the genome data from 22 species of filarial nematode (Table 1). Note that we used the published genome assembly of *Dr. repens* (GCA_008729115.1; [60]) as it had much greater contiguity and higher completeness than either of the two short-read assemblies we produced in assembling the *Wolbachia* genomes for this species.

*Wolbachia* genome sequences were downloaded from NCBI Genomes and complemented with the assembled *Dr. repens Wolbachia* genomes (Table S2). *Wolbachia* is robustly placed within the Rickettsiales in Alphaproteobacteria, and previous work has identified species in *Anaplasma*, *Ehrlichia* and the newly described Candidatus *Mesenet* as its closest relatives. We added the genomes of two *Mesenet* isolates (*Mesenet* endosymbiont of *Phosphuga atrata*, GCA_964020175.1, and *Mesenet* endosymbiont of *Agriotes lineatus*, GCA_964019585.1), two *Ehrlichia* species (*Ehrlichia chaffeensis*, CP000236, and *Ehrlichia ruminantium*, CR925677) and two *Anaplasma* species (*Anaplasma centrale*, CP001759, and *Anaplasma marginale*, CP001079) as the outgroup. The dataset of 1,444 *Wolbachia* genomes is largely composed of genomes from supergroups A and B, with good representation of filarial-nematode-infecting strains. Supergroup C *Wolbachia* were from the following hosts: *Dr. immitis* [38], *O. gibsoni* [61], *O. gutturosa* [62], *O. ochengi* [63] and *O. volvulus* (GCA_000530755.1). Supergroup D *Wolbachia* were included for the following hosts: *B. malayi* [64], *B. pahangi* [65], *Li. brasiliensis* [38], *Li. sigmodontis* [66] and *W. bancrofti* [67]. Supergroup F *Wolbachia* were included from nematode hosts *Ma. perstans* [67], *Ma. ozzardi* [16] and *Md. hiepei* [38], and from the arthropod hosts *Cimex lectularius* [38,68], *Ctenocephalides felis* [8], *Melophagus ovinus* [16], *Menacanthus eurysternus* [69], *Osmia caerulescens* [16] and *Penenirmus auritus* [69]. Supergroup J *Wolbachia* from *Cr. tuberocauda* [64], *Dp. caudispina* [64] and *Dp. gracile* [70] were included.

Completeness of the *Wolbachia* genomes was assessed using CheckM [71] (version 1.2.2). We deployed dRep dereplicate [37] (version 3.4.3) to reduce the redundancy in the dataset, yielding a representative dataset of 165 genomes (see Table S2). A database of these 165 *Wolbachia* genomes was built with kraken [72] (version 2.0.7) and all nematode nuclear genomes were screened for similarity to *Wolbachia*. All flagged contigs and scaffolds were manually assessed with NCBI BLAST+ [73] to detect all those having similarity to *Wolbachia* over the full length of the contig. We decided to remove these sequences from the nuclear assemblies as they most likely derived from living *Wolbachia* rather than nuclear insertions. A list of discarded sequences can be found in Table S3.

### Nematode phylogenetic analysis

BUSCO [74] (version 5.4.7) was run on the genomes of 22 filarial nematodes to detect conserved single-copy genes using the nematode gene set (Nematoda_odb10). Retaining only those genes present across all genomes yielded a set of 670 protein-coding genes. These protein sequences were aligned using MAFFT [40] (version 7.490) in automatic mode. These alignments (a total of 409,399 positions) were analysed with IQ-TREE [44] (version 2.1.4) using protein model GTR20+G4 with 1,000 bootstraps. The phylogeny was rooted with *S. labiatopapillosa*.

### *Wolbachia* phylogenetic analysis

We re-predicted proteomes for the 165 selected *Wolbachia* genomes with prokka [35] (version 1.14.6), which uses prodigal [75] (version 2.6.3) for gene finding. While this automated procedure may introduce errors, especially in prediction of proteins from pseudogenes, prodigal is a robust and accurate gene finder and the kinds of errors it may introduce were unlikely to affect later stages of our analyses. In particular, strain-specific mispredictions will be classified as strain-unique singletons, and excluded from subsequent analyses as uninformative. Orthologues were predicted across these proteomes using OrthoFinder [36] (version 2.4.0). Orthologous families were reduced to only retain dRep-selected genomes. We selected the 696 clusters where at least 75% of the 165 *Wolbachia* strains were single-copy. The protein alignments for these genes were aligned with MAFFT (version 7.490) and trimmed of ambiguous regions using trimAl [76] (version 1.4; -gt 0.25 -st 0.001). The trimmed alignments were concatenated into an alignment of 216,966 amino acid positions and analysed with IQ-TREE (version 2.2.5) using protein model GTR20+G4 with 1000 bootstraps. Additionally, a maximum likelihood phylogenetic tree for each of the 696 genes was inferred with IQ-TREE (version 2.1.4; iqtree -s {alignment} -nt {threads}). Coalescent analysis of these gene trees was performed using ASTRAL [77] (version 5.7.4). The gene-wise coalescent tree was compared to the concatenated sequence tree and no differences in tree topology were observed.

### Construction of hidden Markov models and screening of nuclear genomes for NUWTs

Protein sequences of the 2,647 orthologous groups that had three or more members across more than three *Wolbachia* strains were aligned using MAFFT (version 7.490). The untrimmed MAFFT protein alignment for each cluster was transformed to a nucleotide alignment of the source *Wolbachia* genomic sequences using tranalign from EMBOSS [41] (version 6.6). Each alignment was used to derive a nucleotide hidden Markov model (HMM) in hmmbuild from the hmmer suite [43] (version 3.3.1). The set of 2,647 HMMs was screened against each of the cleaned target nematode genomes using nhmmscan. The tabular output format of nhmmscan was parsed using a python script (SelectMatches.py) to identify putative NUWTs with E-values less than 1e-30. Matches were found for 44.5% (1,179) of the HMMs. We recognise that the nematode genome assemblies have been generated from different data types, are of different qualities (from highly fragmented to chromosomally-scaffolded) and have undergone different contamination purging processes, and thus that the NUWT repertoire of the lower assembly quality genomes may be incomplete. For the highly fragmented assemblies, we removed from the NUWT catalogue any small contigs that had identity to the cognate *Wolbachia* genomes.

The coordinates of the matches were used to direct extraction of the corresponding span of nucleotide sequence from the nuclear genomes (FetchRegionSeq.py). These nucleotide sequences were profile aligned to the cognate tranalign alignments using MAFFT. Where the resulting alignment contained more than fifty supergroup A or supergroup B *Wolbachia* sequences, only dRep selected genomes were retained for these supergroups (using python script Reduce_alignment.py). The alignment was used to position the putative NUWTs within the diversity of the sequences from living *Wolbachia* using IQ-TREE (version 2.1.4) with a substitution model search (-m MFP) [78].

To identify the likely supergroup-of-origin of the NUWTs, the 1,179 output trees were traversed with a python script (Reroot_rename_tree.py) that uses the ete package from etetoolkit [79] (version 3.1.2). Trees were first rooted, either using outgroup sequences or, if these were not present, by midpoint rooting. All *Wolbachia* sequences were renamed according to their supergroup classification and nodes that had a uniform classification were collapsed. The placement of all putative NUWTs in their trees was assessed and NUWTs were classified according to the supergroup of their closest sister branch.

NUWT locations were converted to bed files and NUWTs were clustered into regions using bedtools [80] (version 2.29.0) if the next NUWT was located within 2.5 kb and there were no intervening coding genes.

### Nematode gene and repeat annotation

For all 22 nematode genomes we performed *de novo* repeat identification using EarlGrey [81] (version 3.0). We provided the curated repeat libraries to RepeatMasker (https://www.repeatmasker.org/; version 4.1.5) to soft-mask transposable elements. Protein-coding annotation for the filarial nematodes were taken from the following sources: Wormbase Parasite [82] for *A. viteae*, *B. pahangi*, *B. timori*, *Ce. johnstoni*, *Dr. immitis*, *E. elaphii*, *Lo. loa, O. flexuosa*, *O. ochengi* and *W. bancrofti*, a public Braker annotation [18] for *Cr. tuberocauda*, *Dp. caudispina*, *Li. brasiliensis*, *Md. hiepei* and *Ma. ozzardi* and an Augustus annotation [18] for *M. perstans*. Liftoff [83] (version 1.6.3) was used for *A. viteae*, *B. pahangi*, *Ma. perstans* and *W. bancrofti* to transfer the annotation to the genome version used in this study. For the four species without public annotation; *Dr. repens*, *Li. sigmodontis*, *O. lupi* and *S. labiatopapillosa,* BRAKER [84] (version 3) was run on soft-masked genomes using only protein alignments from related species as evidence by using nematoda OrthoDB [85] (version 11), which is comprised of 147,547 protein sequences.

Enrichment for overlap of NUWTs with different repeat categories was calculated using regioneR [86] (version 3.20), which performs a permutation test (10,000 times) while maintaining chromosome structure. Results were afterwards corrected for multiple testing with Benjamini-Hochberg.

### Data availability

New nematode genome sequences are available in INSDC. New *Wolbachia* genome sequences are available in INSDC. A list of *Wolbachia* genome sequences analysed is available in supplementary tables. The Supplementary Data, Figures and Tables have been uploaded to Zenodo under doi 10.5281/zenodo.15032066.

## Supplementary Materials

### Supplementary data

Supplementary data files are available at Zenodo under doi **10.5281/zenodo.15032066**

The zip archive includes three directories:

### Scripts

Python scripts written for data processing.

FetchRegionSeq.py.gz

Reduce_alignment.py.gz

Reroot_rename_tree.py.gz

SelectMatches.py.gz

### Data

Directories containing intermediate data files generated for the analyses presented.

*1_WolbachiaAnnotation* - the prokka annotation of each *Wolbachia* genome analysed

*2_WolbachiaOrthoFinder* - the orthoFinder results for the clustering of the proteomes of the 165 selected *Wolbachia* proteomes

*3_WolbachiaPhylogeny* - the phylogeny of *Wolbachia* inferred using Astral

*4_WolbachiaOrthoAlignments* - protein alignments of orthogroups from the orthoFinder analysis of 165 selected *Wolbachia* peoteomes

*5_WolbachiaHMM* - nucleotide hidden Markov models generated from the nucleotide alignments of orthogroups from the orthoFinder analysis of 165 selected *Wolbachia* proteomes

*6_NUWTtrees* - phylogenies inferred using Astral of the *Wolbachia* orthogroups for which NUWT sequences were identified in the filarial nematode genomes. The NUWT sequences were profile aligned to the orthogroup nucleotide alignments.

*7_NUWTmatches* - files giving the nucleotide coordinates and scores of each orthogroup HMM in the filarial genomes

### Supplementary Tables

(in Excel format)

Table S1: *Wolbachia* genome data

Table S2: Dereplicated *Wolbachia* genomes

Table S3: Contigs removed from filarial nematode genomes as likely contaminants.

Table S4: Numbers and classification of NUWTs detected in each filarial nematode species

**Figure S1:**
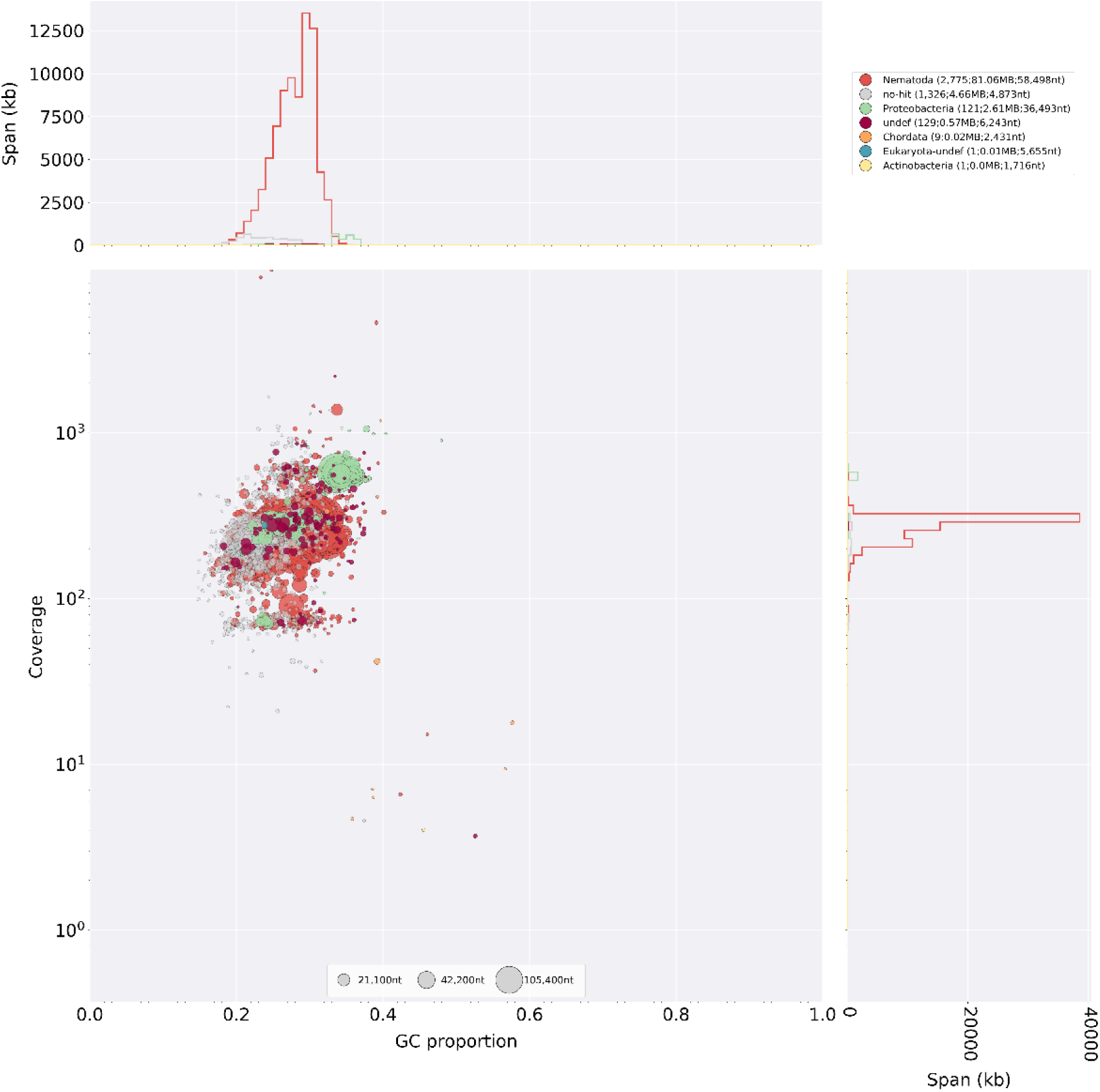
BlobTools plot of initial *Dirofilaria repens* assembly. A BlobTools (v1) plot of the *Dr. repens* assembly showing (red) contigs assigned to Nematoda (i.e. the nematode host) and (green) contigs assigned to Proteobacteria (i.e. *Wolbachia*). The Protobacteria contigs (coloured green) form two clusters, one at ∼300 fold coverage and 35% GC, and one at ∼200 fold coverage and 28% GC. These correspond to the C supergroup (high coverage) and F supergroup (low coverage) *Wolbachia* coinfecting this sample.

**Figure S2:**
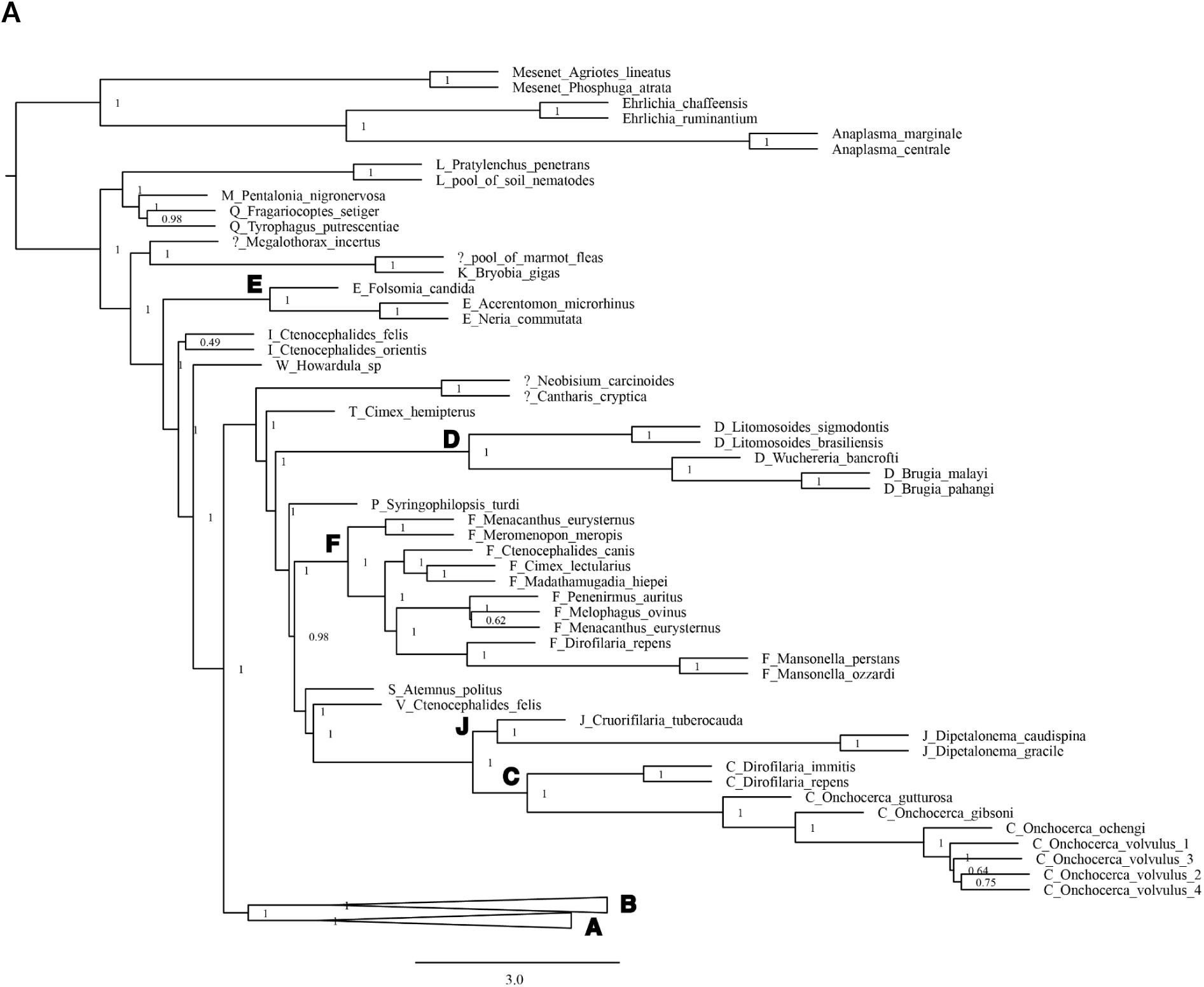
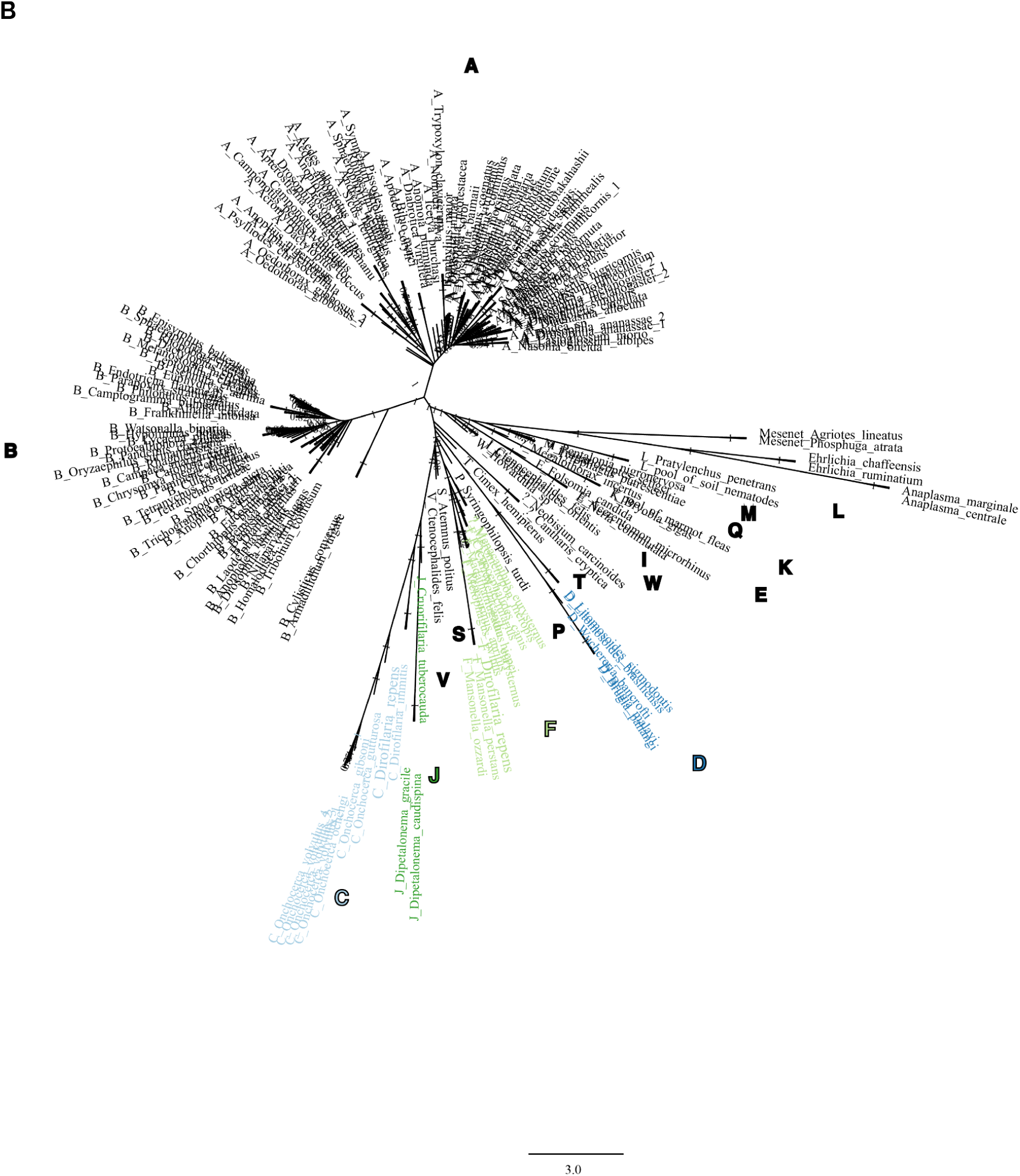
Genome phylogeny of *Wolbachia*. **A.** Summary phylogeny of 1,444 *Wolbachia* genomes. The phylogeny was rooted with genomes from seven *Anaplasma*, *Ehrlichia* and *Mesenet* species. Supergroups are indicated with bold letters. **B.** Phylogeny of 167 selected *Wolbachia* genomes (as in Figure 1B) with sources named. Supergroups are indicated with bold letters.

**Figure S3:**
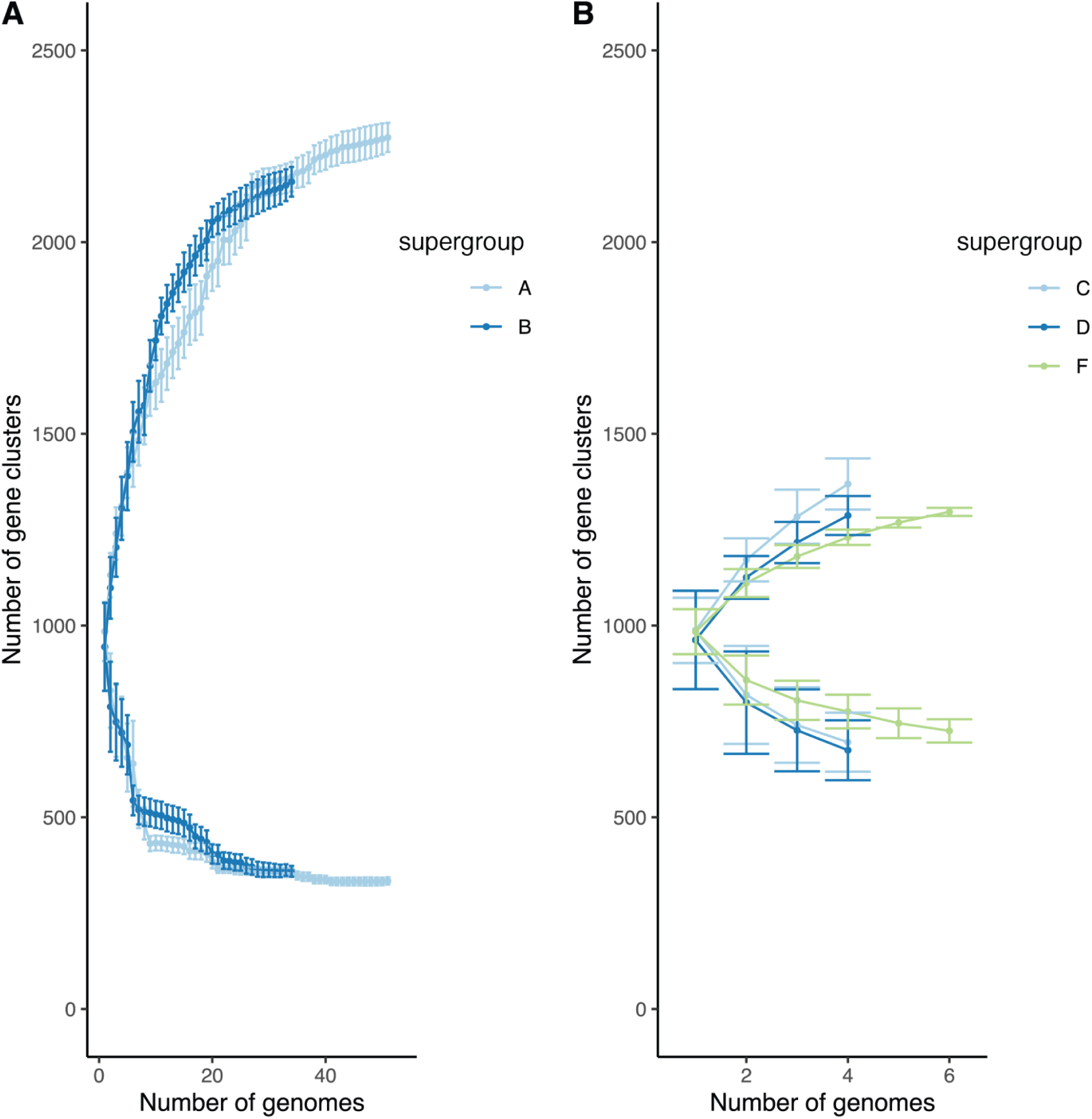
*Wolbachia* proteome clustering. **A.** Rarefaction curves describing the proteome diversity found across *Wolbachia* and within supergroup A and B. *Wolbachia* proteomes selected with dRep and deemed near-complete were added one by one and the number of clusters with >1 member tallied. The standard deviations are derived from 50,000 repetitions of the clustering with randomised addition order of genomes. The upper curves are for orthogroups with >1 member, while the lower curves indicate the number of singleton sequences. **B.** Rarefaction curves describing the proteome diversity found within supergroup C, D and F. *Wolbachia* proteomes selected with dRep and deemed near-complete were added one by one and the number of clusters with >1 member tallied. The standard deviations are derived from 50,000 repetitions of the clustering with randomised addition order of genomes. The upper curves are for orthogroups with >1 member, while the lower curves indicate the number of singleton sequences.

**Figure S4:**
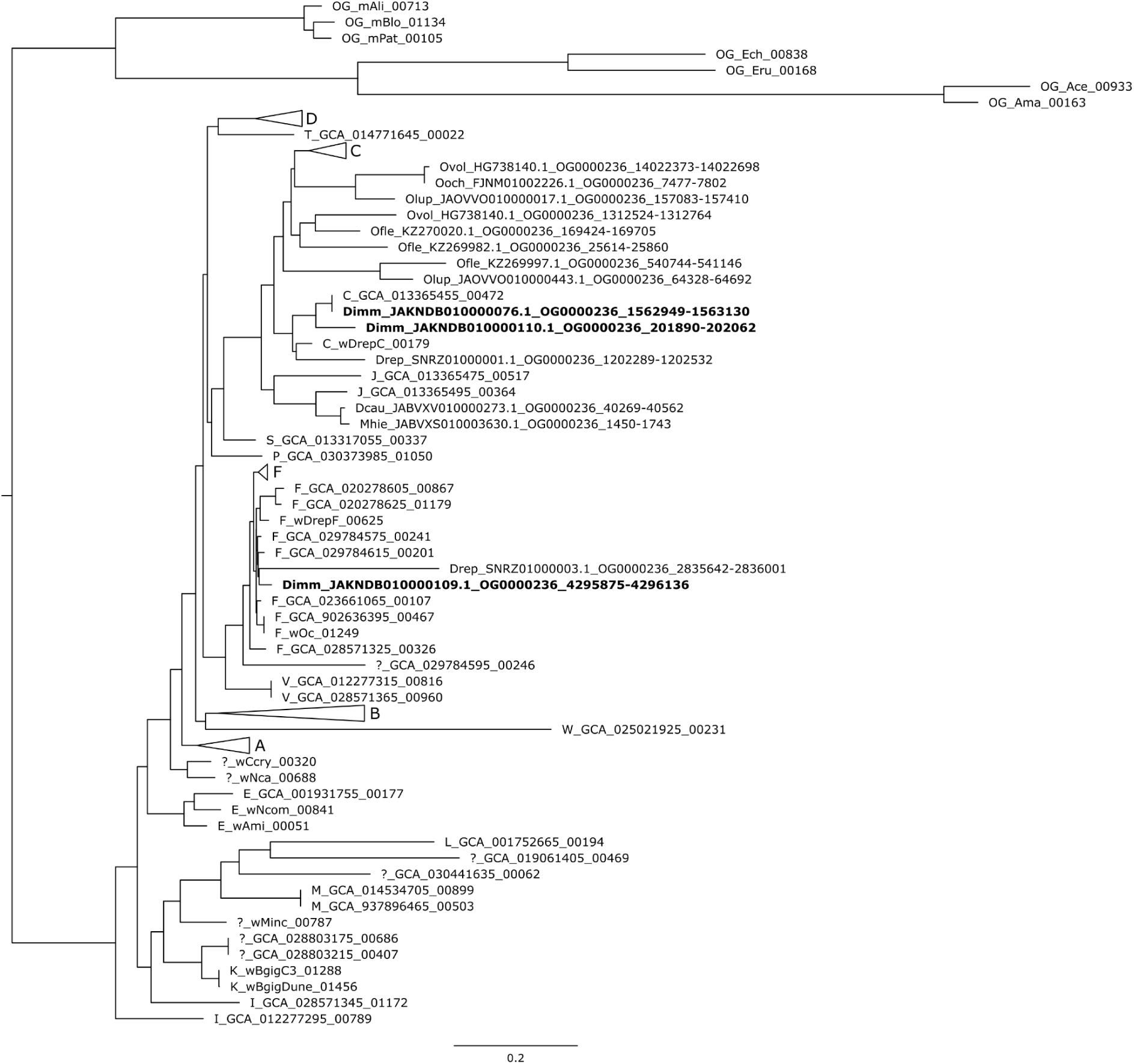
Phylogenetic tree of orthologous family OG0000236 and NUWTs. Phylogenetic tree of dihydrolipoyl dehydrogenase (OG0000236), illustrating the phylogenetic placement of NUWTs deriving from both C and F *Wolbachia* in *Dirofilaria immitis* (highlighted in bold). Sequences derived from living *Wolbachia* are indicated by the supergroup and host species nomenclature followed by the locus number (e.g. “E_wNcom_00841”) or supergroup and GCA nomenclature followed by the locus number (e.g. “I_GCA012277295_00789”), while NUWTS are indicated by the nematode species abbreviation and their location (e.g. “Dimm_JAKNDB010000109.1_OG0000236_4295875-4296316” is from *D. immitis*, contig JAKNDB010000109.1; it matches OG0000236 and is from bases 4295875-4296316 in the contig).

**Figure S5:**
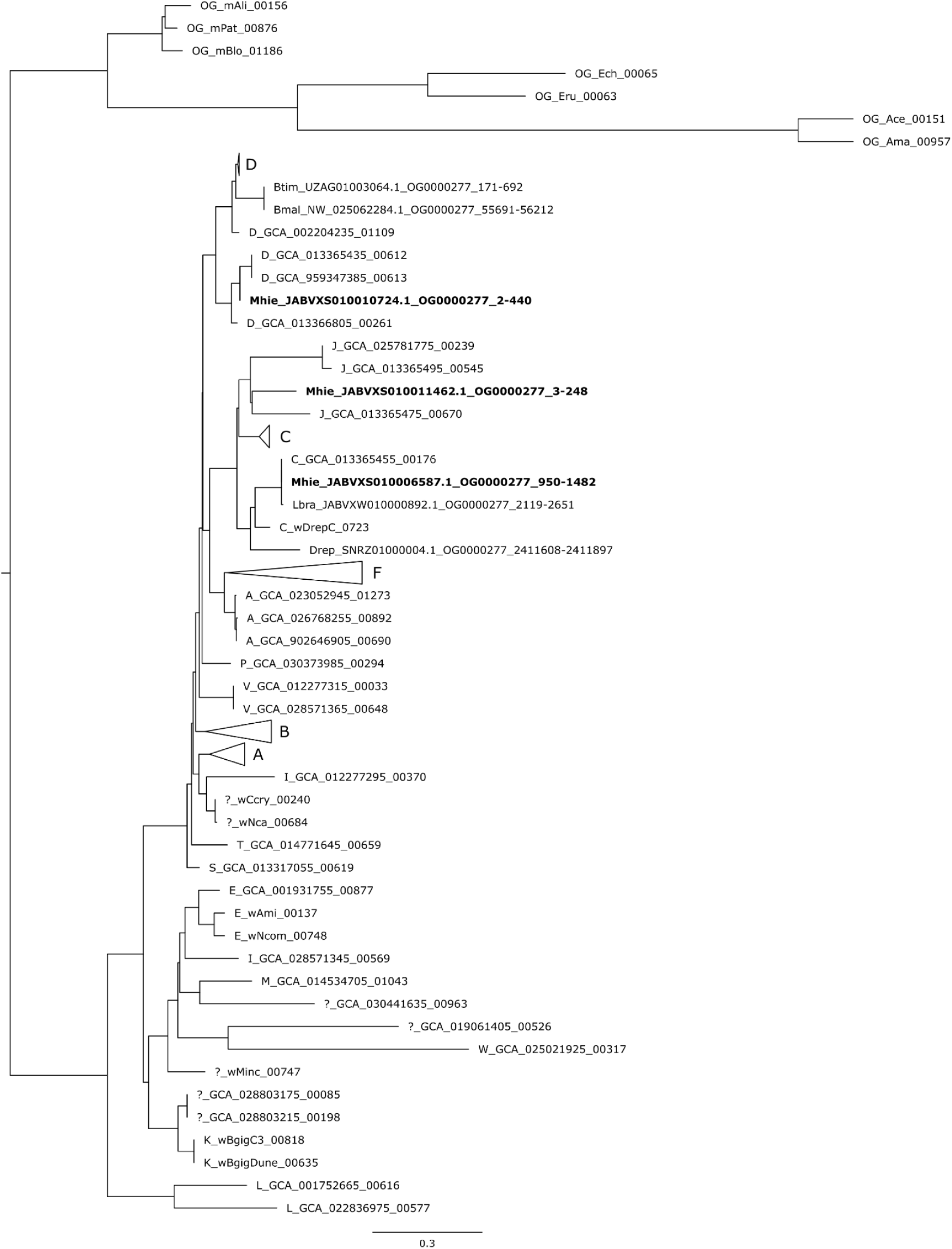
Phylogenetic tree of orthologous family OG0000277 and NUWTS. Phylogenetic tree of peptide deformylase (OG0000277) showing the phylogenetic placement of NUWTs from *Madathamugadia hiepeia*. The NUWTs likely derive from C, D and J *Wolbachia*. Nomenclature as in Figure S4.

**Figure S6:**
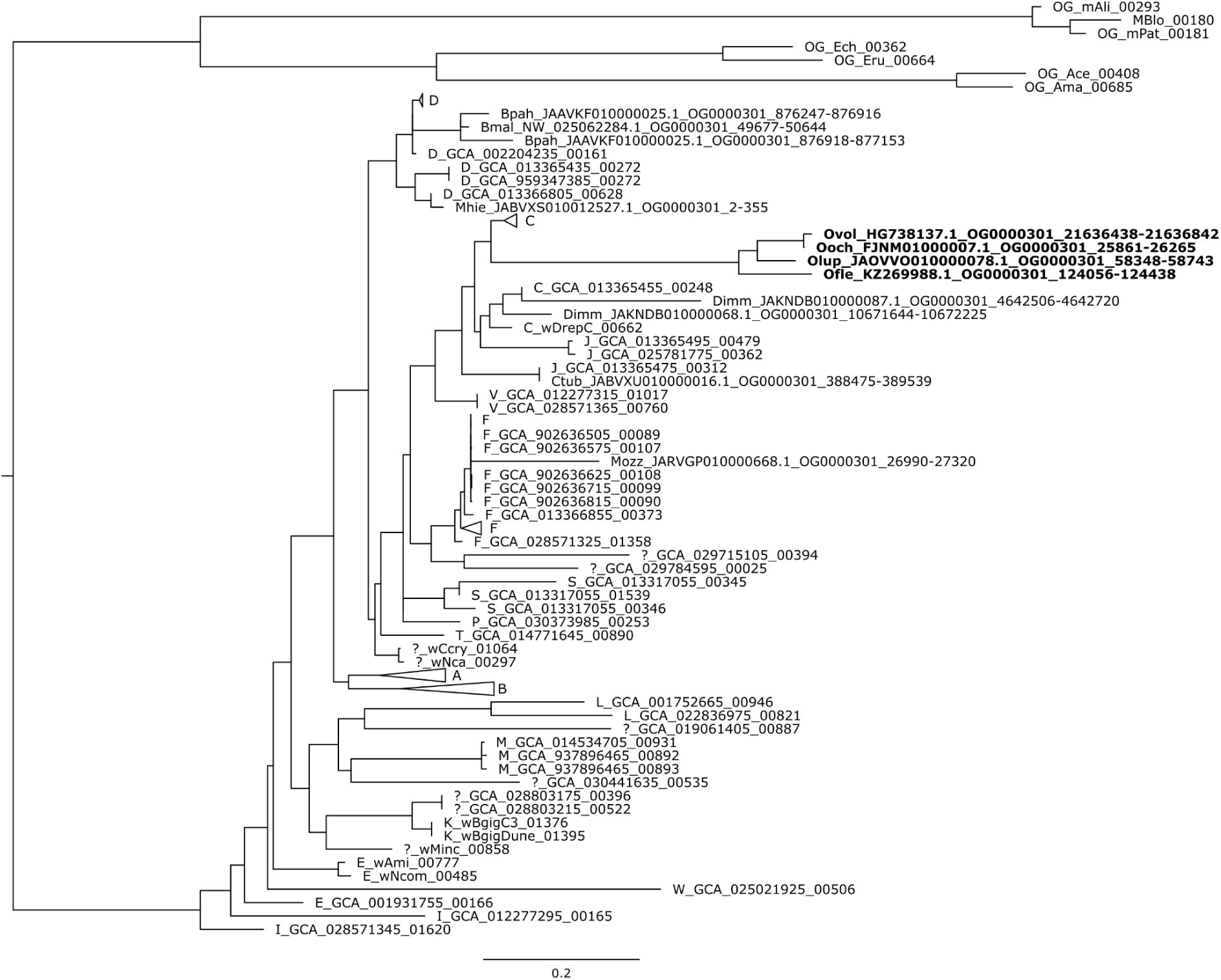
Phylogenetic tree of orthologous family OG0000301 and NUWTs. Phylogenetic tree of DNA-directed RNA polymerase subunit alpha (OG0000301) showing the phylogenetic placement of NUWTs from four *Onchocerca* species. Nomenclature as in Figure S4.

**Figure S7:**
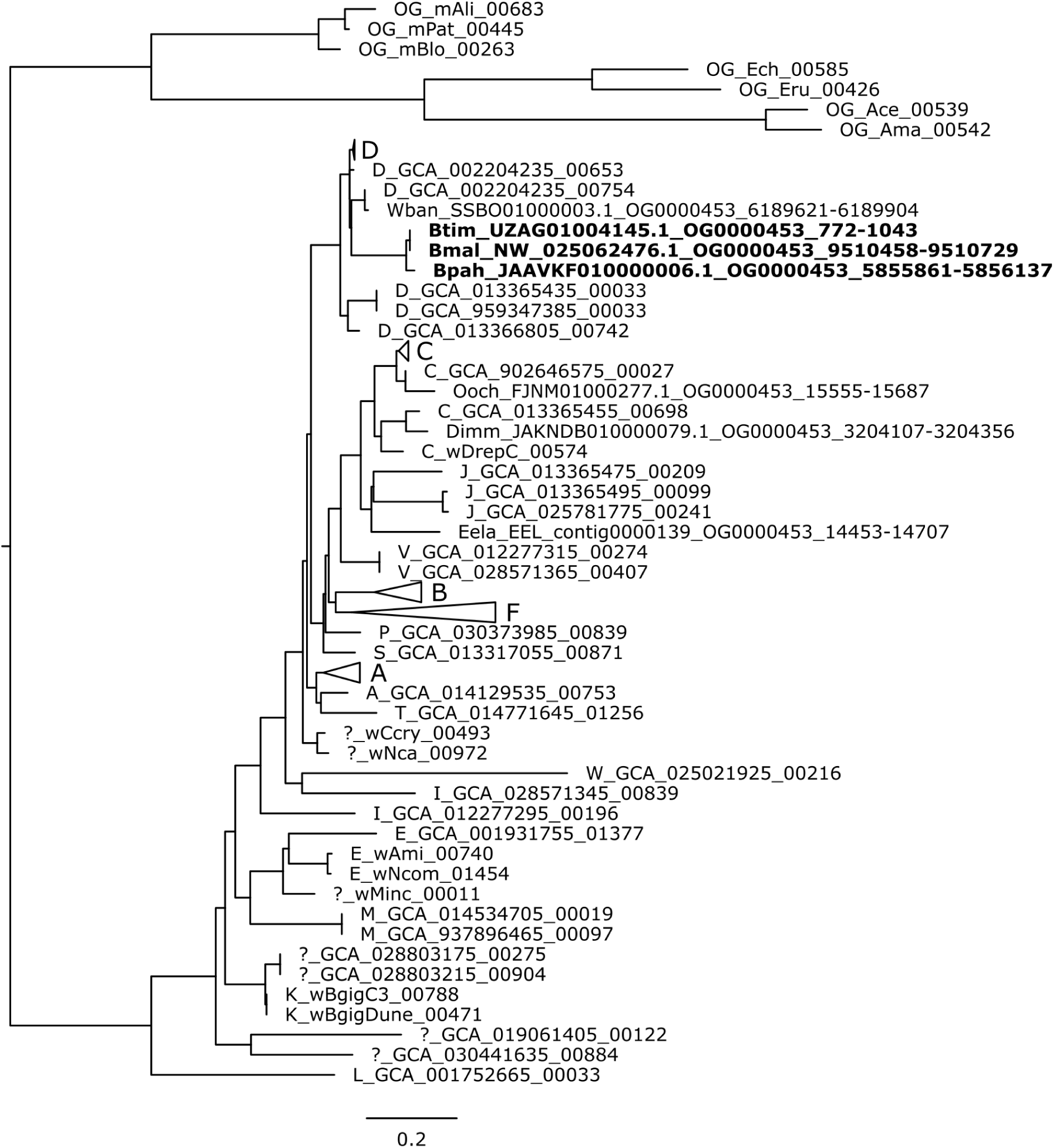
Phylogenetic tree of orthologous family OG0000453 and NUWTs. Phylogenetic tree of signal peptidase I (OG0000453) showing the phylogenetic placement of NUWTs from three *Brugia* species. Nomenclature as in Figure S4.

**Figure S8:**
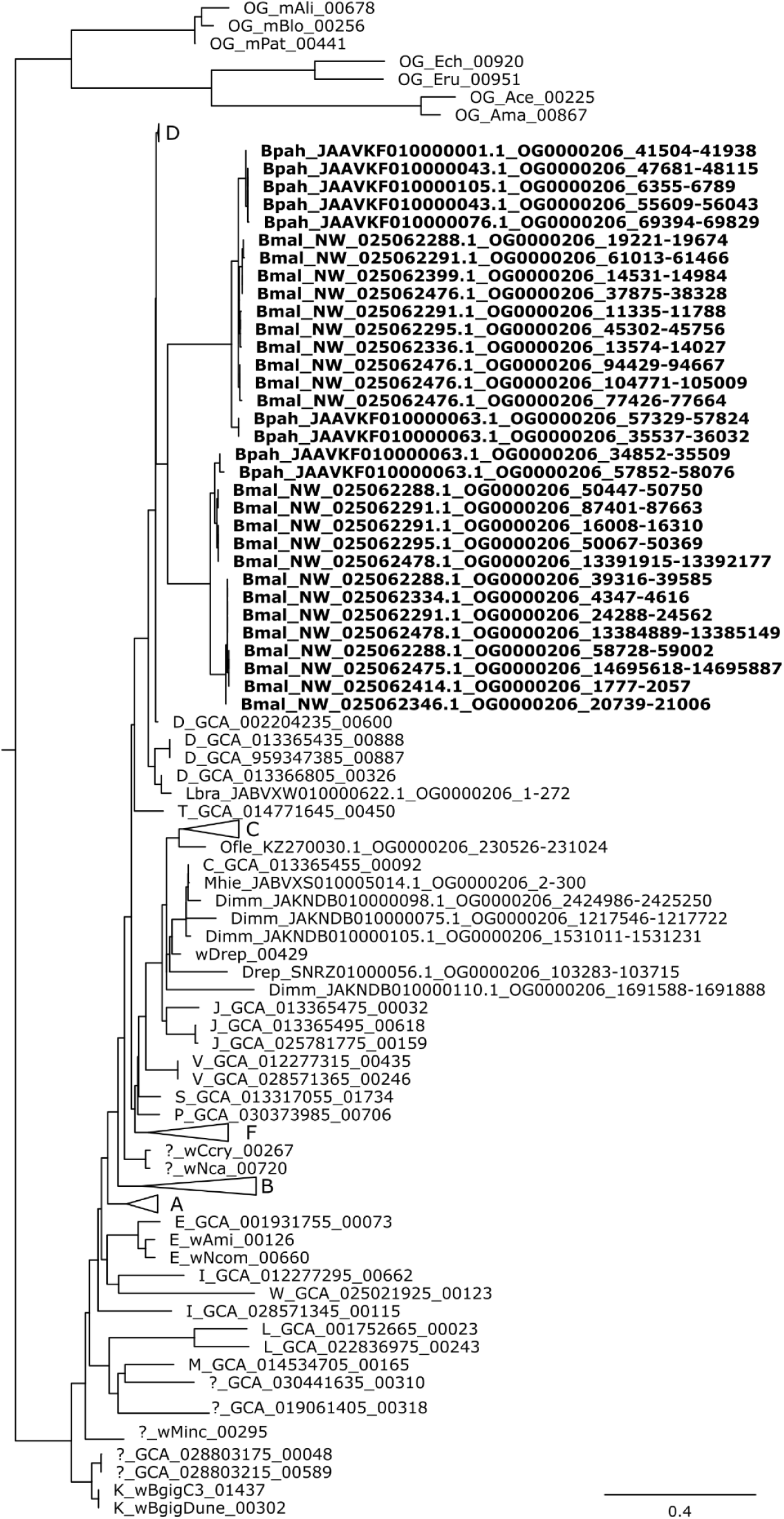
Phylogenetic tree of orthologous family OG0000206 and NUWTs. Phylogenetic tree of ATP-dependent zinc metalloprotease FtsH (OG0000206), showing the phylogenetic placement of multi-copy NUWTs from two *Brugia* species. Nomenclature as in Figure S4.

